# Impairments in the mechanical effectiveness of reactive balance control strategies during walking in people post-stroke

**DOI:** 10.1101/2022.07.28.499225

**Authors:** Chang Liu, Jill L. McNitt-Gray, James Finley

**Affiliations:** Department of Biomedical Engineering, University of Southern California, Los Angeles, USA; Department of Biological Science, University of Southern California, Los Angeles, USA; Division of Biokinesiology and Physical Therapy, University of Southern California, Los Angeles, USA; Neuroscience Graduate Program, University of Southern California, Los Angeles, CA, 90089

**Keywords:** balance, stroke, angular impulse, angular momentum, gait, reactive control, forward loss of balance

## Abstract

People post-stroke have an increased risk of falls compared to neurotypical individuals, partly resulting from an inability to generate appropriate reactions to restore balance. However, few studies investigated the effect of paretic deficits on the mechanics of reactive control strategies following forward losses of balance during walking. Here, we characterized the biomechanical consequences of reactive control strategies following perturbations induced by the treadmill belt accelerations. Thirty-eight post-stroke participants and thirteen age-matched and speed-matched neurotypical participants walked on a dual-belt treadmill while receiving perturbations that induced a forward loss of balance. We computed whole-body angular momentum and angular impulse using segment kinematics and reaction forces to quantify the effect of impulse generation by both the leading and trailing limbs in response to perturbations in the sagittal plane. We found that perturbations to the paretic limb led to larger increases in forward angular momentum during the perturbation step than perturbations to the non-paretic limb or to neurotypical individuals. To recover from the forward loss of balance, neurotypical individuals coordinated reaction forces generated by both legs to decrease the forward angular impulse relative to the pre-perturbation step. They first decreased the forward pitch angular impulse during the perturbation step. Then, during the first recovery step, they increased the backward angular impulse by the leading limb and decreased the forward angular impulse by the trailing limb. In contrast to neurotypical participants, people post-stroke did not reduce the forward angular impulse generated by the stance limb during the perturbed step. They also did not increase leading limb angular impulse or decrease the forward trailing limb angular impulse using their paretic limb during the first recovery step. Lastly, post-stroke individuals who scored poorer on clinical assessments of balance and had greater motor impairment made less use of the paretic limb to reduce forward momentum. Overall, these results suggest that paretic deficits limit the ability to recover from forward loss of balance. Future perturbation-based balance training targeting reactive stepping response in stroke populations may benefit from improving the ability to modulate paretic ground reaction forces to better control whole-body dynamics.

## 1 Introduction

People post-stroke have an increased risk of falls relative to neurotypical individuals (Weerdesteyn et al., 2008) and this may be due, in part, to impairments in their ability to generate appropriate reactive strategies following a loss of balance. These impairments result from a combination of delayed muscle activation to external perturbations (Kirker et al., 2000; Marigold et al., 2004), abnormal muscle activation patterns (Higginson et al., 2006), and weakness (Olney & Richards, 1996). In addition, trips or slips, which commonly occur in the direction of walking, are one of the most prevalent causes of falls among people post-stroke (Schmid et al., 2013). Although prior studies have examined the dynamics of backward losses of balance during stance (Patel & Bhatt, 2017; Salot et al., 2016) and walking post-stroke (Dusane et al., 2021; Kajrolkar et al., 2014; Kajrolkar & Bhatt, 2016), few have investigated the mechanics and recovery strategies following forward losses of balance during walking.

When responding to forward losses of balance during walking, neurotypical individuals adopt a sequence of reactive control strategies across multiple steps to counteract the forward rotation of the body (Debelle et al., 2020). For example, when people trip over an obstacle, their first opportunity to recover balance involves modulating the support limb’s push-off force to reduce forward angular momentum (Pijnappels et al., 2005). Next, people often increase the length of the recovery step to reduce forward momentum while walking (Debelle et al., 2020; Golyski et al., 2022; Mathiyakom & McNitt-Gray, 2008; Roeles et al., 2018; Vlutters et al., 2016). As a result, the ground reaction forces of the leading recovery limb and the perturbed trailing limb combine to generate a backward moment about the center of mass (CoM) and help arrest the forward rotation of the body (Mathiyakom & McNitt-Gray, 2008).

However, sensorimotor deficits in people post-stroke may prevent them from executing successful reactions to forward losses of balance while walking. If a perturbation occurs during paretic stance, the paretic leg may be too weak to adequately support the body or it may lack the dexterity to properly regulate the body’s momentum (Arene & Hidler, 2009; Honda et al., 2019; Nott et al., 2014; Roerdink et al., 2009). Therefore, people post-stroke may not have sufficient time to step further forward with the non-paretic limb and arrest forward momentum. Conversely, if a perturbation occurs during non-paretic stance, they may have difficulty initiating a successful stepping response with the paretic leg to help restore balance due to deficits in paretic propulsion (Allen et al., 2014; Chen et al., 2005; Lauzière et al., 2015) and hip flexion (Rybar et al., 2014). However, it has yet to be determined how paretic deficits impact the biomechanical consequences of reactive response to forward losses of balance or whether these effects differ following perturbations to the paretic versus non-paretic limbs.

Here, our objective was to determine how stroke influences the biomechanical consequences of reactive control strategies following sudden treadmill accelerations (Figure 1). To counteract the increase in forward angular momentum following a perturbation, participants could use a combination of recovery strategies during the perturbation and recovery steps. First, they could reduce the forward angular impulse during the single stance phase following the perturbation. Second, they could increase the backward angular impulse generated by the leading limb during the recovery step. Finally, they could also decrease the forward angular impulse generated by the trailing limb during the first recovery step. We hypothesized that treadmill accelerations would cause larger increases in forward angular momentum in people post-stroke compared to neurotypical control individuals regardless of the side of the perturbation as post-stroke deficits may prevent these individuals from generating adequate reactive control strategies. We also expected that perturbations of the paretic leg would lead to greater increases in forward angular momentum than perturbations of the non-paretic side due to deficits in the ability of the paretic leg to support body weight (Figure 1A). When considering the biomechanical consequences of the reactive responses, we hypothesized that neurotypical participants would have larger contributions to the reduction of forward angular momentum from both the perturbed limb and the recovery limb compared with those of people post-stroke (Figure 1B-C). Lastly, we hypothesized that post-stroke participants would generate smaller reductions in forward angular impulse by the perturbed limb and larger increases in backward angular impulse using the recovery limb during the first recovery step following paretic versus non-paretic perturbations (Figure 1B-C).

**Figure 1:**
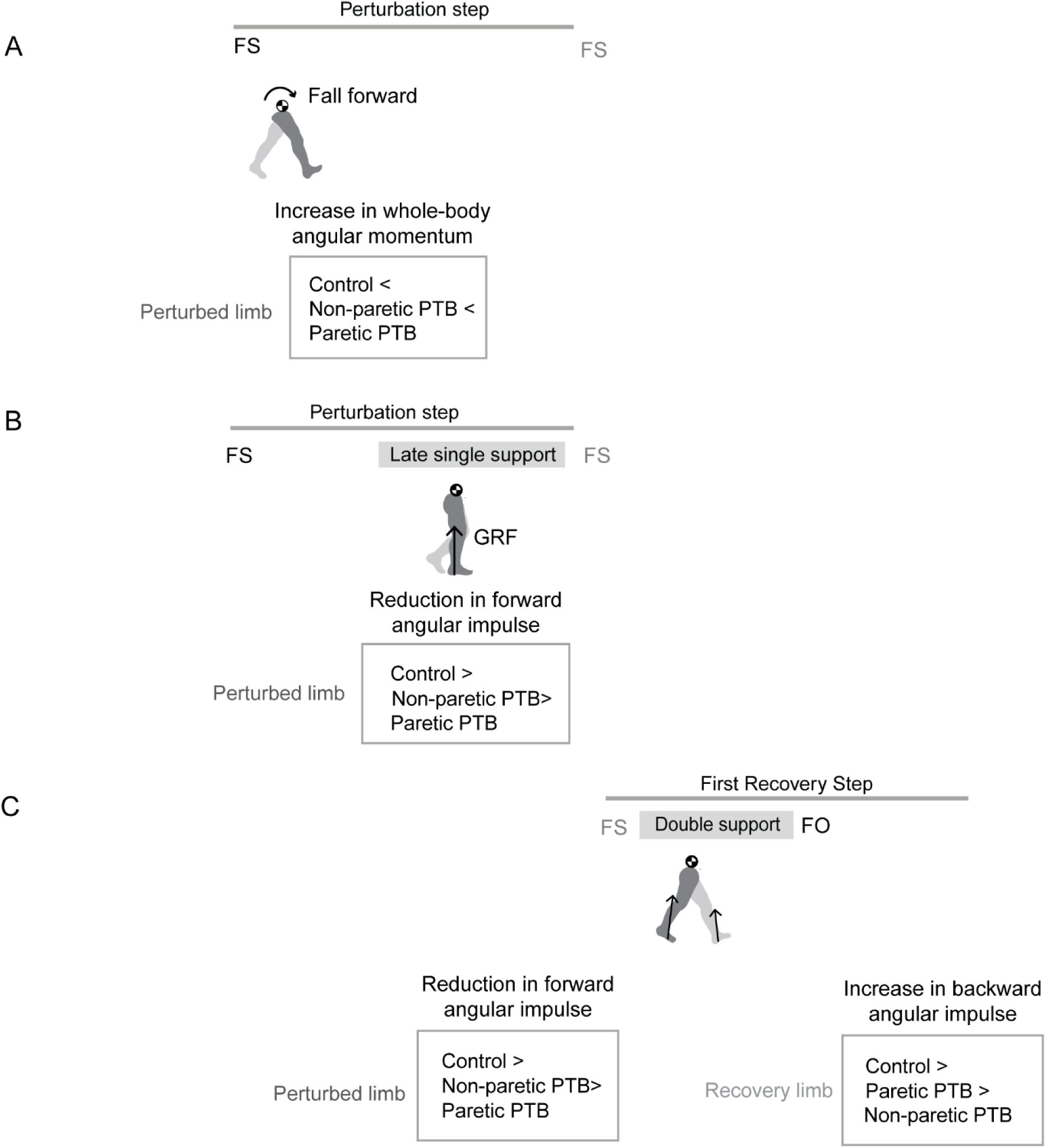
Graphical illustration of hypothesized differences in whole-body angular momentum and angular impulses. (A) Hypothesis for changes in whole-body angular momentum during the perturbation step and the first recovery step relative to that measured during the pre-perturbation step. (B) Hypothesis for changes in angular impulse during the late single support phase of the perturbation step relative to that measured during the pre-perturbation step. (C) Hypothesis for changes in the trailing perturbed limb angular impulse and the leading limb angular impulse during the double support phase of the first recovery step relative to that measured during the pre-perturbation step. (FS: foot strike; FO: foot-off; GRF: ground reaction force; PTB: perturbation)

## 2 Methods

### 2.1 Participants

We recruited 38 people post-stroke (Table 1) from the IRB-approved, USC Registry for Aging and Rehabilitation, the USC Physical Therapy Associates Clinic, and Rancho Los Amigos National Rehabilitation Center. Inclusion criteria for the stroke survivors were the following: 1) a unilateral brain lesion 2) paresis confined to one side, 3) ability to walk on the treadmill for five minutes without holding on to any support. Use of ankle-foot orthoses was permitted during the experiment. We also recruited 13 age-matched neurotypical participants from the community. Exclusion criteria for neurotypical participants were neurological, cardiovascular, orthopedic, and psychiatric diagnoses. Study procedures were approved by the Institutional Review Board at the University of Southern California and all participants provided written, informed consent before testing began. All aspects of the study conformed to the principles described in the Declaration of Helsinki.

**Table 1.** Participant demographics for both control and stroke participants. Values are formatted as Mean (SD).

|  | **Control**<br>**(N = 13)** | **Stroke**<br>**(N= 38)** | **p value** |
| --- | --- | --- | --- |
| Age (yrs) | 58 (29) | 60 (11) | 0.76 |
| Female/Male | 6/7 | 14/24 | / |
| Mass (kg) | 76 (15) | 81 (19) | 0.38 |
| Height of CoM (m) | 0.97 (0.05) | 0.94 (0.06) | 0.13 |
| Treadmill speed (m/s) | Matched: 0.6 (0.2) | 0.6 (0.2) | 0.62 |
| Scaling factor $\sqrt{gL}$ (m/s) | 3.08 (0.082) | 3.04 (0.095) | 0.13 |
| Self-selected Overground speed (m/s) | 1.3 (0.2) | 0.8 (0.3) | <0.0001 |
| Berg Balance Scale | 55 (2) | 51 (6) | 0.017 |
| Activity-specific Balance Confidence Scale | 97 (3.5) | 77 (13) | <0.0001 |
| Falls Efficacy Scale | 18 (2) | 29 (12) | 0.0025 |
| Lower Extremity Fugl-Meyer | / | 26 (5) | / |
| Left/right hemiparetic | / | 15/23 | / |
| Months after stroke | / | 83 (55) | / |
| Functional Gait Assessment | / | 21 (6) | / |

### 2.2 Experimental protocol

The experimental protocol for post-stroke participants has been described previously (Buurke et al., 2020), and we provide a summary of the procedures and setup below. The complete protocol consisted of a set of clinical assessments and walking trials on the treadmill. Before the walking trials, we evaluated motor impairment using the lower extremity portion of the Fugl-Meyer Assessment (FM) (Fugl-Meyer et al., 1975), static balance using Berg Balance Scale (BBS)(Berg et al., 1992), static and dynamic balance during locomotion using the Functional Gait Assessment (FGA)(Leddy et al., 2011), and over-ground walking speed using the 10-meter walking test. Participants also completed questionnaires about balance confidence using the Activity-Specific Balance Confidence Scale (ABC) (Powell & Myers, 1995). Higher scores on all these assessments indicated better balance control or higher balance confidence. Lastly, we completed a Fall History Questionnaire for participants who experienced at least one fall within the past year. After clinical evaluations, we instructed stroke participants to walk on the dual-belt treadmill (Bertec, Columbus, OH, USA). A harness was provided to prevent the participants from falling but no body weight support was provided. First, the participants walked on the treadmill to familiarize themselves with the experimental setup. To identify participants’ preferred walking speed on the treadmill, we started from 70% of the speed obtained from a 10-meter walking test and adjusted their walking speed by 0.05 m/s increments or decrements until the participants verbally indicated that they achieved their preferred walking speed (Park et al., 2021). Participants then walked for three minutes at their self-selected speed. After the unperturbed walking trial, participants completed a familiarization trial with at least two sudden treadmill accelerations which were triggered at foot-strike based on the ground reaction forces recorded by the treadmill’s force plates. Finally, participants completed two trials of three minutes at their self-selected speed during which they received six accelerations to the treadmill belts on each side.

Neurotypical participants also completed a set of clinical assessments including the ABC, FES, BBS, and 10-meter walking test. We instructed the participants to walk at matched speeds with a stroke participant of similar age, and they completed one unperturbed walking trial and one perturbed trial at this speed. For the perturbed trial, 10 perturbations occurred on each side.

For both groups, treadmill accelerations were triggered at random intervals within 15 to 25 steps after the previous perturbation to allow participants to reestablish their walking patterns. Each perturbation was characterized by a trapezoidal speed profile in which the speed increased by 0.2 m/s at an acceleration of 3 m/s^2^, was held for 0.7 s, and then decelerated back to the self-selected speed during the swing phase of the perturbed leg. Between each trial, stroke participants had breaks of at least three minutes to minimize fatigue while control participants were given breaks as needed. Participants did not hold on to handrails while walking on the treadmill.

### 2.3 Data Acquisition

A ten-camera motion capture system (Qualisys AB, Gothenburg, Sweden) recorded 3D marker kinematics at 100 Hz and ground reaction forces at 1000 Hz. We placed a set of 14 mm spherical markers on anatomical landmarks (Havens et al., 2018; Song et al., 2012) and placed marker clusters on the upper arms, forearms, thighs, shanks, and the back of heels. Marker positions were calibrated during a five-second standing trial at the beginning of each trial. We removed all joint markers after the calibration.

### 2.4 Data Processing

We post-processed the kinematic and kinetic data in Visual3D (C-Motion, Rockville, MD, USA) and Matlab 2020b (Mathworks, USA) to compute variables of interest. Marker positions and ground reaction forces were low-pass filtered by 4^th^ order Butterworth filters with cutoff frequencies of 6 Hz and 20 Hz, respectively based on previous literature (Kurz et al., 2012; Reisman et al., 2009; Winter, 2009). We defined foot strike as the point when the vertical ground reaction force became greater than 150N and foot off as the point when vertical ground reaction force became less than 150N (Liu et al., 2018). We removed the perturbations that occurred more than ~150 ms after foot strike. We included a median of 11 (interquartile range: 3.5) perturbations per side for each stroke participant and a median of 10 (interquartile range: 0.5) perturbations per side for each age-matched control participant. We categorized the pre-perturbation steps as the last two steps before the perturbation occurred (Pre-PTB_1-2_), perturbation steps (PTB) as the step during which the perturbation was applied, and recovery steps (R_1-3_) as the three steps that followed the perturbation.

### 2.5 Whole-body angular momentum

We created a 13-segment, whole-body model in Visual3D and calculated the angular momentum of each segment about the body’s center of mass for neurotypical participants (Herr & Popovic, 2008; Martelli et al., 2013). The model included the following segments: head, thorax, pelvis, upper arms, forearms, thighs, shanks, and feet. We modeled the limb segments’ mass based on anthropometric tables (Dempster, 1955), and the segment geometry based on the description in Hanavan (Hanavan, 1964). For stroke participants, the pelvis segment was modeled to be rigidly connected to the trunk because they wore an extra harness that blocked the markers necessary to track the pelvis accurately. Sagittal plane angular momentum was defined as the projection of angular momentum on the mediolateral axis passing through the body CoM (Silverman & Neptune, 2011). Whole-body angular momentum (*L*) was computed as the sum of all segmental angular momenta which were composed of segmental rotation about the body’s CoM and rotation of each segment about its CoM. *L* was nondimensionalized by a combination of the participant’s mass (*M*), the participant’s height of COM (*H*), and gravity constant (g) to reduce between-subject variability (Eqn. 1)(Martelli et al., 2013).

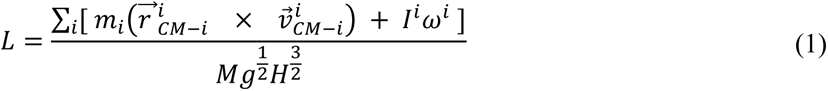

Here, *m* is segmental mass, *r* is the distance from segment to the body COM, *I* is the segmental moment of inertia, ω is the segmental angular velocity, and the index *i* corresponds to individual limb segments. Negative values of angular momentum represented forward rotation, while positive values represented backward rotation. Although we used a 12-segment instead of a 13-segment model for people post-stroke, this had a negligible effect on whole-body angular momentum. The root-mean-square error for the peak backward and forward whole-body angular momentum in the sagittal plane between the 12-segment model and the 13-segment model was 2.1 ± 1.5 % and 0.95 ± 0.70 %, respectively (Park et al., 2021). We computed integrated whole-body angular momentum (L_int_) for each step cycle to characterize changes in the body configuration over each step (Liu et al., 2018; Potocanac et al., 2014).

### 2.6 Measures of reactive control strategies

In addition to whole-body angular momentum, we used angular impulse to quantify the mechanical consequences of the reactive control strategies on whole-body dynamics. We determined the effect of the ground reaction forces from each limb on the change in whole-body dynamics using measures of angular impulse as described in Eqn.2-3 (Figure 2). Similar to whole-body angular momentum, sagittal plane angular impulse was defined as the projection of angular impulse on the mediolateral axis passing through the body CoM. The earliest strategy that people could employ to begin recovering from losses of balance during the perturbation step is to modulate the ground reaction force of the perturbed limb to reduce the forward momentum about the CoM. Such a strategy is expected to occur no less than ~200ms after the onset of a perturbation and this would approach the late single-support phase of the gait cycle (Sloot et al., 2015). Thus, we first computed the forward pitch impulse (ΔL_Stance_) during the late single support phase to capture the effect of the perturbed stance limb ground reaction force on whole-body dynamics (Eqn.2). The forward pitch impulse is mathematically equivalent to the change in whole-body angular momentum during the single support phase.

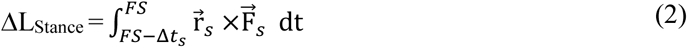

**Figure 2:**
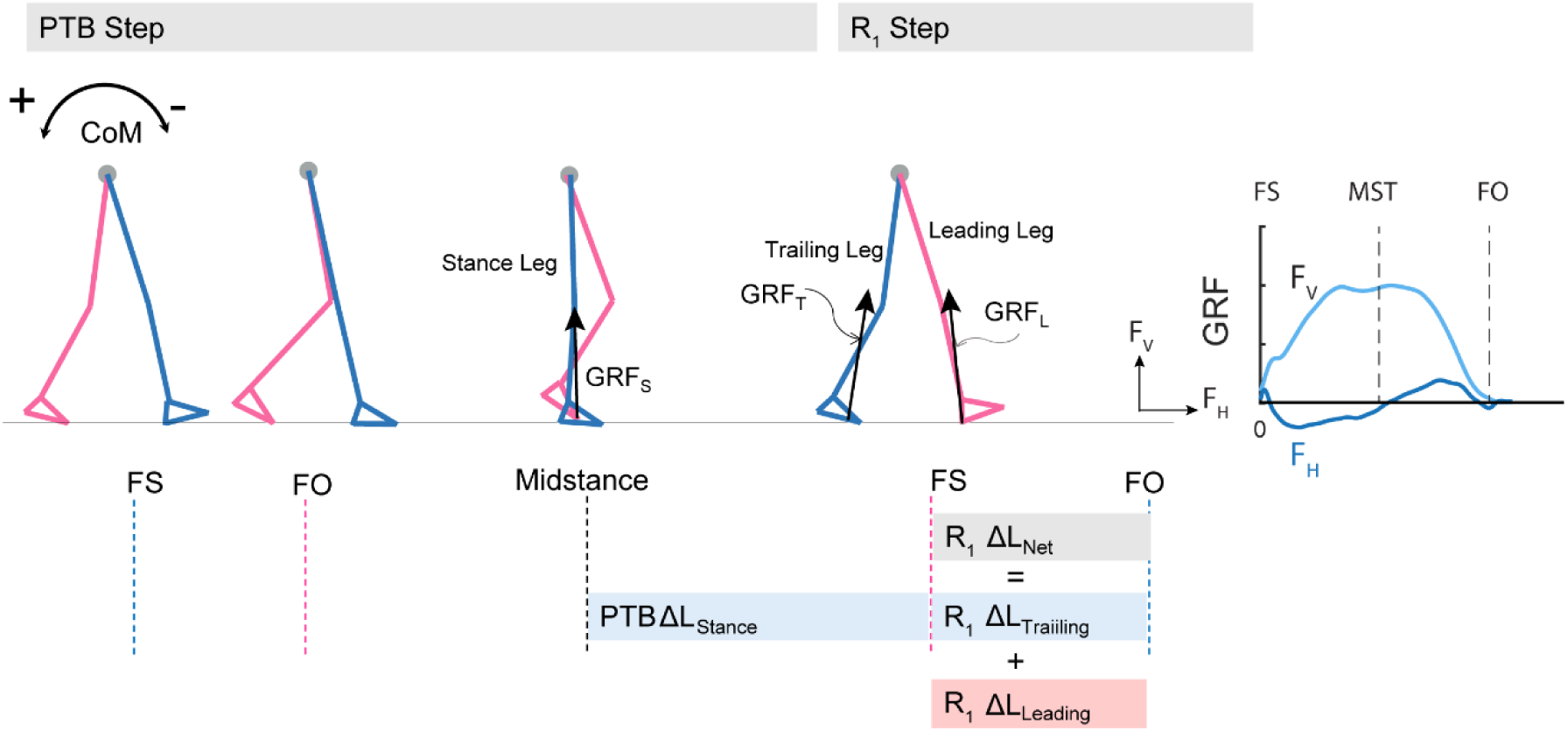
Diagram of computed angular impulse about the body CoM by the leading and trailing leg during the perturbation (PTB) step and the first recovery step (R1) and illustration of ground reaction force from foot-strike to foot-off during one example perturbation step. The forward pitch impulse (PTB ΔL_Stance_) is computed during the phase from midstance of the PTB step until the foot strike of the R_1_ step. Net angular impulse (R_1_ ΔL_Net_) is computed as the sum of trailing limb angular impulse (R_1_ ΔL_Trailing_) and leading limb angular impulse (R_1_ ΔL_Leading_) during the double support phase of the R_1_ step. FS: Foot strike; FO: Foot-off. The arrows (+/-) indicate the backward and forward moments by the GRF about CoM, respectively. F_v_: vertical ground reaction force; F_H_: fore-aft ground reaction force.

Here, 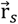 represents the displacement vector from the body’s CoM to the center of pressure of the stance limb. 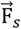 represents the stance limb’s ground reaction force. We defined the duration of the late single support phase (Δ*t_s_*) as 80% of the average time from midstance to the subsequent foot strike during pre-perturbation steps. Midstance was defined as the midpoint between consecutive foot strikes during pre-perturbation steps. We used the same time duration across all step types to remove the effect of time on computing angular impulses. Index *s* corresponds to the stance leg.

We also computed the net angular impulse (ΔL_Net_) during the double support phase of the recovery step as the sum of the leading limb (ΔL_Leading_) and trailing limb (ΔL_Trailing_) angular impulse (Eqn.3). The contributions to the net angular impulse from the leading and trailing limbs were computed similar to Eqn. 2, except that the integration was performed from foot-strike (FS) to FS + Δ*t_ds_* Δ*t_ds_* represents the double support phase (Adamczyk & Kuo, 2009). We again used 80% of the average double support time during pre-perturbation steps so that the same duration was used across all step types.

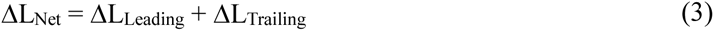

### 2.7 Statistical Analysis

All statistical analyses were performed in Matlab 2020b (Mathworks). For people post-stroke, if the non-paretic leg was perturbed, the Pre-PTB step, PTP step, R_2_, and R_4_ steps were non-paretic steps, and the R_1_ and R_3_ steps were paretic steps, and vice versa for the paretic perturbations.

We first tested whether there were significant differences in any participant characteristics between control and post-stroke participants by using a two-sample t-test with unequal variances. We also tested whether there were significant differences in L_int_, ΔL_stance,_ ΔL_Net_, ΔL_Leading_, and ΔL_Trailing_ during the pre-perturbation step between the control and stroke group (paretic and non-paretic steps) and within the stroke group (paretic vs. non-paretic steps). We analyzed the normality of these measures using the Shapiro-Wilk Test. We used a two-sample unequal variance t-test if the data were normally distributed; otherwise, we used the Mann-Whitney test. We adjusted for multiple comparisons using Bonferroni corrections.

We then assessed if any of the dependent variables L_int_, ΔL_Net_, ΔL_Leading_, and ΔL_Trailing_ following perturbations differed from those measured during the pre-perturbation step using linear mixed-effect models for stroke participants and control participants, respectively. The independent variables for this analysis included Step Type (Pre-PTB_1-2_, PTB, R_1-3_), side of perturbation (Leg) (paretic and non-paretic side), and the interaction between Step Type and Leg to determine if changes in any of the dependent variables from the pre-perturbation step differed between sides. The reference level was set to be Pre-PTB_1_. For neurotypical participants, we did not find that any of the variables differed between sides. Thus, we combined values across limbs for the remainder of the analysis and the independent variable only included Step Type (Pre-PTB_1-2_, PTB, R_1-3_). We included a random intercept for each model to account for unmodeled sources of between-subject variability. We also determined the number of recovery steps needed for participants to restore balance by identifying when *L_int_* returned to values measured before the perturbations. We analyzed the angular impulse during the perturbation and first recovery steps as our prior work demonstrated that reactive stabilization strategies were most evident during these two steps (Liu et al., 2018). We used the Shapiro-Wilk Test to test the residual normality. We provide detailed statistical results in Table S1 for this analysis.

We also determined if the deviation of the dependent variables from pre-perturbation values differed between neurotypical participants and stroke participants following paretic and non-paretic perturbations. We used a two-sample unequal variance t-test if the variables were normally distributed; otherwise, we used the Mann-Whitney test, and the comparisons between groups were adjusted for multiple comparisons using Bonferroni corrections. We provided detailed statistical results for this test of normality in Table S2. Lastly, we computed Pearson correlation coefficients to test for associations between changes in ΔL_stance_ during the perturbation step, changes in ΔL_Leading_ and ΔL_Trailing_ during the first recovery step relative to the pre-perturbation step, and each clinical balance assessment (BBS, FGA, ABC, FM). Significance was set at α = 0.05.

## 3 Results

### 3.1 Whole-body angular momentum

The acceleration of the belts caused consistent increases in forward angular momentum and triggered multi-step balance recovery responses for both neurotypical participants and people post-stroke (Figure 3, first row). During the perturbation step, angular momentum became more negative as the body rotated forward. To compensate for the perturbation, participants then generated positive angular momentum and initiated backward rotation during the first recovery step (Figure 3, first row). We also computed the integrated angular momentum over each step to characterize changes in body configuration in response to perturbations. Participants increased their forward rotation, indicated by a more negative *L_int_,* during the perturbation step relative to the pre-perturbation step (Figure 4). They then countered the effects of the perturbation during the first recovery step (R_1_) as indicated by a more positive *L_int_*. Neurotypical participants restored whole-body angular momentum to levels comparable to those observed during the pre-perturbation step by the second recovery step while people post-stroke restored angular momentum to pre-perturbation values by the third recovery step (Figure 4).

**Figure 3:**
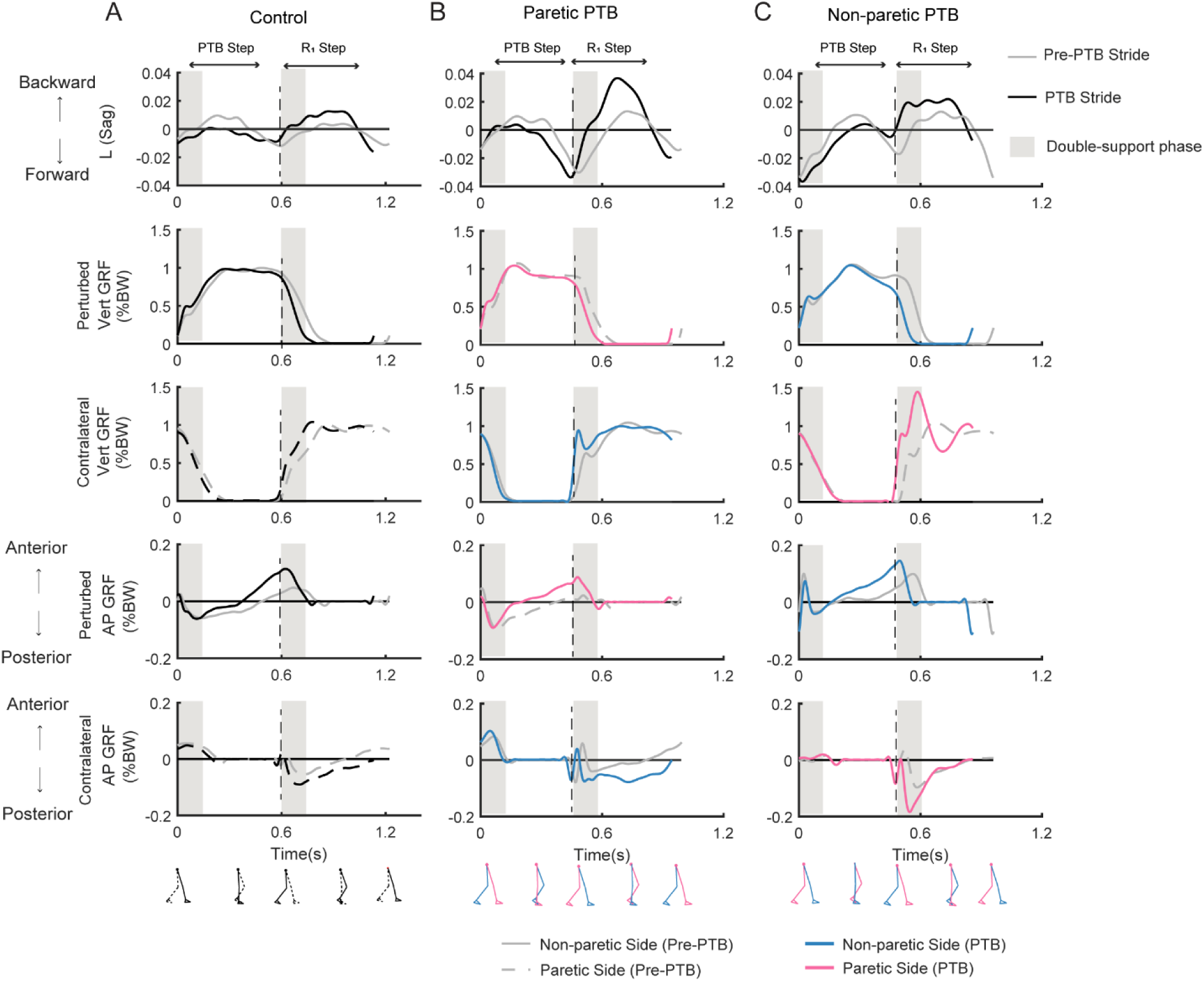
Whole-body angular momentum in the sagittal plane and ground reaction forces for one representative neurotypical participant (A) and a stroke participant during a paretic perturbation (B) and non-paretic perturbation (C) for both a pre-perturbation stride and a perturbation stride. Each stride began at foot strike. The gray traces indicate the time series data for a pre-perturbation stride while the black or colored traces indicate a perturbation stride. Negative values of angular momentum represent forward rotation while positive values represent backward rotation. Ground reaction forces (% body weight) in the vertical and anterior-posterior directions for the perturbed and the contralateral limb when perturbations occurred on the dominant side for the neurotypical participant (A), or on the paretic (B) or non-paretic sides (C) for the stroke participant. For the neurotypical participant, black lines indicated the perturbed side, and the dashed lines indicated the contralateral side. For the stroke participant, pink and blue lines represent the paretic leg and non-paretic leg, respectively. Black dashed vertical lines correspond to the time of foot strike. Gray shaded vertical box corresponds to the double support phase from the time of foot strike to the contralateral foot-off. Pre-PTB: pre-perturbation stride; PTB: perturbation stride.

**Figure 4.**
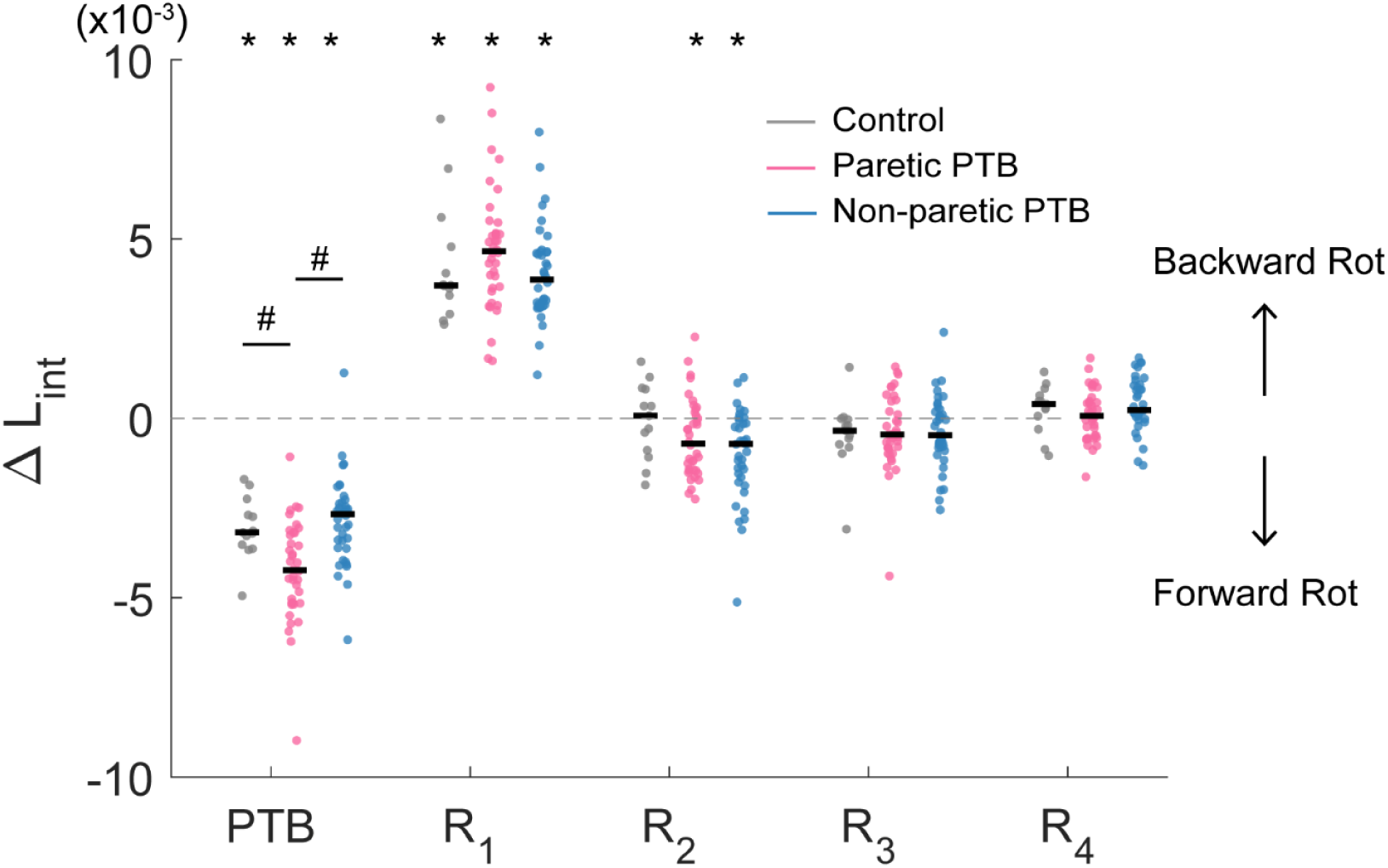
Median integrated angular momentum in the sagittal plane over the step cycle relative to the corresponding pre-perturbation step (ΔL_int_) for all participants (N = 38 stroke participants and N = 13 neurotypical participants). Each dot represents one participant. Black horizontal lines indicate the median across participants. Steps alternated between paretic and non-paretic for stroke participants. PTB: Perturbation step; R: Recovery step. The asterisks on top of the boxplots indicate whether the difference in L_int_ from the pre-perturbation step was significantly different from zero (*p < 0.05) and the # indicated that the ΔL_int_ was different between groups. Note that for people post-stroke, if the non-paretic leg was perturbed, the R_1_ steps were paretic, and Pre-PTB steps and PTB steps were non-paretic and vice versa for the paretic perturbations.

Stroke participants (36 out of 38) increased integrated angular momentum more during paretic perturbations relative to non-paretic perturbations (p = 0.021), indicating that they fell forward more when the perturbation occurred during paretic stance. The increase in integrated angular momentum during the perturbation step was higher during paretic perturbations than for neurotypical participants (Bonferroni corrected p = 0.018), but there was no difference in the increase in integrated angular momentum between non-paretic perturbations and those for neurotypical participants (p = 0.56).

### 3.2 Changes in the stance-phase forward pitch impulse during the perturbation (ΔL_Stance_)

We did not observe any difference increase in forward pitch impulse between neurotypical participants and stroke participants during the perturbation step (Bonferroni corrected p>0.05) or any difference between limbs in people post-stroke (p = 0.088). The earliest strategy that people could employ to begin recovering from losses of balance during the perturbation step is to modulate the ground reaction force of the perturbed limb to reduce the forward angular momentum about CoM. The ground reaction force produced by the stance limb from midstance to the subsequent foot strike typically produced a forward pitch impulse (Figure 5A, Figure S1). During the perturbation step, neurotypical participants produced a smaller forward pitch impulse relative to the pre-perturbation step (p = 0.0005, Figure 6) indicating that they began to arrest the forward loss of balance during the perturbation step. However, people post-stroke only decreased the forward pitch impulse during non-paretic perturbations (29 out of 38 participants, p = 0.005) and not during paretic perturbations (p = 0.67, Figure 6). Although 30 of 38 participants had greater reductions in forward pitch impulse during non-paretic perturbations compared to paretic perturbations, there was no significant difference between sides (p = 0.088).

**Figure 5.**
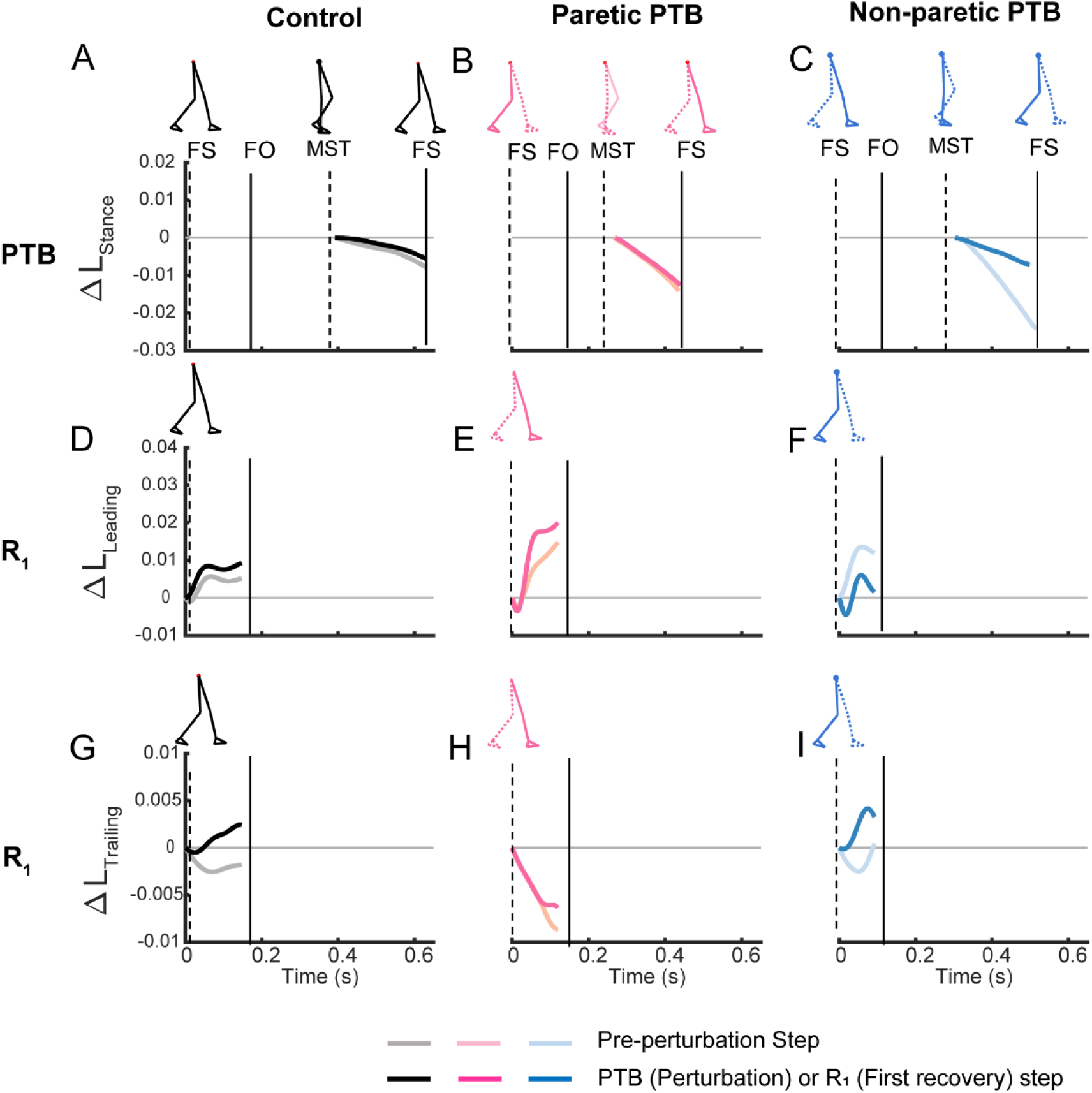
Time series trajectories of the PTB ΔL_Stance_, R_1_ΔL_Leading_, R_1_ΔL_Trailing,_ and the corresponding trajectories during the pre-perturbation step for a representative neurotypical perturbation (left), one paretic perturbation (middle), and one non-paretic perturbation (right). Pre-perturbation trajectories are shown in lighter colors while perturbation and recovery traces are shown in darker colors. The first, second, and third rows correspond to PTB ΔL_Stance_, R_1_ ΔL_Leading_, and R_1_ ΔL_Trailing_, respectively. FS: Foot strike, FO: Foot-off, MST: Midstance. Vertical lines indicate gait events with the solid line corresponding to the ipsilateral limb while the dashed line indicates the contralateral limb.

**Figure 6.**
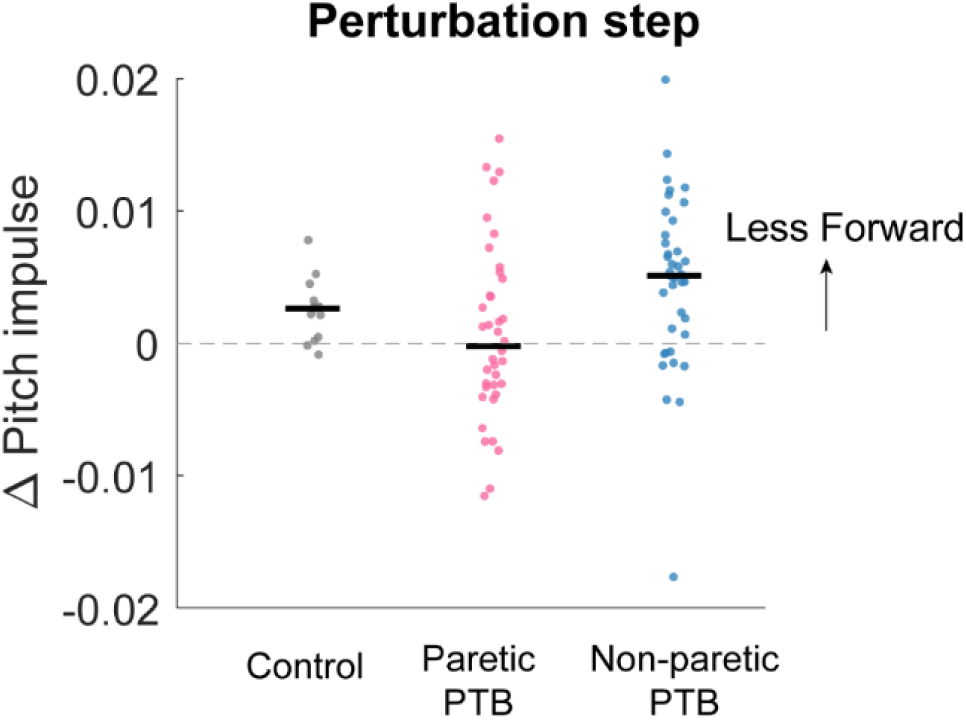
Median changes in pitch impulse following perturbations compared to those measured during the pre-perturbation step for control participants (Gray, N = 13) and during paretic (Pink) and non-paretic (Blue) steps for stroke participants (N=38). Each dot represents one participant. Black horizontal lines indicate the median across participants. Positive values indicate less forward pitch impulse. (B) The asterisks (*) indicate whether the group mean is significantly different from zero (*p<0.05, **p<0.001).

### 3.3 Changes in the net angular impulse during the recovery step (ΔL_Net_)

Neurotypical participants increased net angular impulse from the pre-perturbation step more than stroke participants during the double support phase of the first recovery step following paretic perturbations (Bonferroni corrected p = 0.0072) but this increase did not differ from stroke participants following non-paretic perturbations (Bonferroni corrected p = 0.051). For the pre-perturbation step, the net angular impulse was typically positive during the double support phase for neurotypical individuals, indicating that the ground reaction forces by the leading and trailing limbs generated a net increase in backward angular momentum during this period (Figure S2A). During the first recovery step following a perturbation, neurotypical participants increased the net angular impulse (p<0.0001, Figure 7A), which helped reduce the forward momentum generated by the perturbation. For people post-stroke, the net angular impulse during the first recovery step increased from the pre-perturbation step following non-paretic perturbations (27 out of 38 participants, p = 0.0003) but not following paretic perturbations (p = 0.1, Figure 7A). This result suggests that people post-stroke did not arrest the forward falls as completely when perturbations occurred on the paretic side.

**Figure 7.**
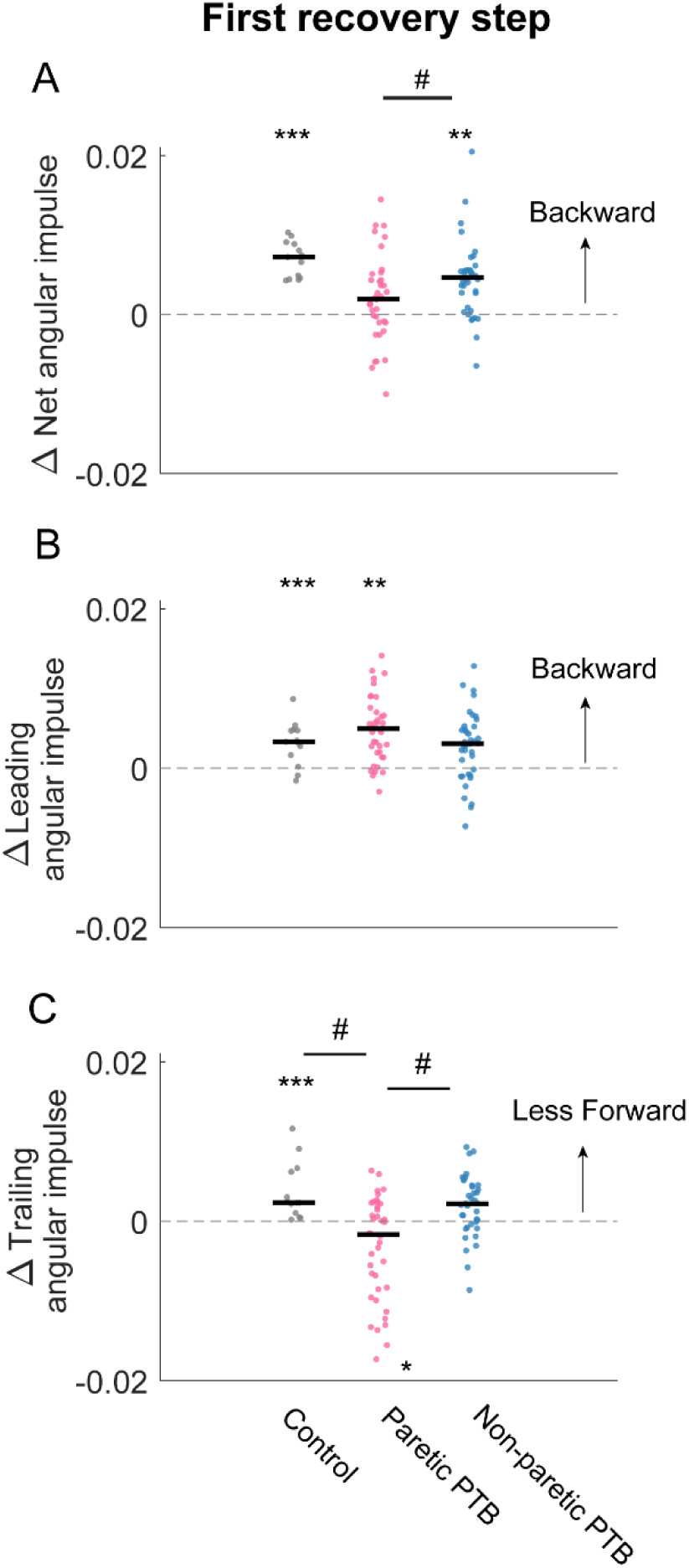
Changes in median net, leading limb, and trailing limb angular impulse during the first recovery step compared to those measured during the pre-perturbation step. Changes in net angular impulse (A), leading limb angular impulse (B), and trailing limb angular impulse (C) during the first recovery step compared to the pre-perturbation step. Each dot represents one participant. Black horizontal lines indicate the median across participants. The asterisks (*) indicate whether values were statistically different from zero (*p<0.05, **p<0.001, ***p<0.0001). The hashes (#) indicate when comparisons between groups are significantly different. Note that for people post-stroke, if the non-paretic leg was perturbed, the leading and trailing limbs corresponded to the paretic and non-paretic limbs during the first recovery step and vice versa for the paretic perturbations.

### 3.4 Changes in the leading limb angular impulse during the recovery step (ΔL_Leading_)

There was no difference in the increase in leading limb backward impulse during the first recovery step between stroke and neurotypical participants (All Bonferroni corrected p>0.05). During the pre-perturbation step, the ground reaction force generated by the leading leg of neurotypical control participants produced a backward angular impulse about the CoM during the double support phase (Figure 5D, Figure S2C). During the double support phase of the first recovery step, neurotypical participants increased this backward impulse to help arrest the forward fall, and this was evidenced by a more positive leading limb angular impulse for neurotypical participants (p < 0.0001, Figure 7B). For stroke participants, leading limb angular impulse increased from the pre-perturbation step following paretic perturbations (33 out of 38 participants, p = 0.0006) but not following non-paretic perturbations (p = 0.075, Figure 7B).

### 3.5 Changes in the trailing limb angular impulse during the recovery step (ΔL_Trailing_)

Neurotypical participants reduced trailing limb angular impulse more than stroke participants following paretic perturbations (Bonferroni corrected p = 0.0033) but not following non-paretic perturbations (Bonferroni corrected p = 0.69). Stroke participants also reduced forward trailing limb angular impulse more by the non-paretic limb following non-paretic perturbations than following paretic perturbations (29 out of 38 participants, p = 0.029). During the pre-perturbation step, the ground reaction force by the trailing limb generated a forward moment about the body’s CoM and thus the trailing limb angular impulse was negative for neurotypical participants (Figure 5G, Figure S2E). Neurotypical participants decreased their forward angular impulse which indicates that the trailing limb assisted with recovery from a forward loss of balance. During the first recovery step, forward trailing limb angular impulse did not change from the pre-perturbation step following non-paretic perturbations (p = 0.27) for stroke participants. However, forward trailing limb angular impulse increased following paretic perturbations from the pre-perturbation step (21 out of 38 participants, p = 0.047).

### 3.6 Association between reactive stabilization strategies and clinical measures

Lastly, we assessed whether changes in forward pitch impulse during the perturbation step, leading limb angular impulse during the first recovery step, and trailing limb angular impulse during the first recovery step from the pre-perturbation steps were associated with clinical assessment of balance and motor impairment (BBS, FGA, ABC, FM) in our sample of stroke participants. We found significant correlations between paretic trailing limb angular impulse and scores on clinical assessments of balance and motor impairment. The reduction in trailing limb angular impulse following the paretic perturbations relative to the pre-perturbation step was positively correlated with FM (R^2^= 0.31, p = 0.0002) and FGA (R^2^= 0.23, p = 0.002, Figure 8). This indicated that participants who scored poorer on clinical assessments of balance and had greater motor impairment made less use of the paretic limb to reduce forward momentum.

**Figure 8:**
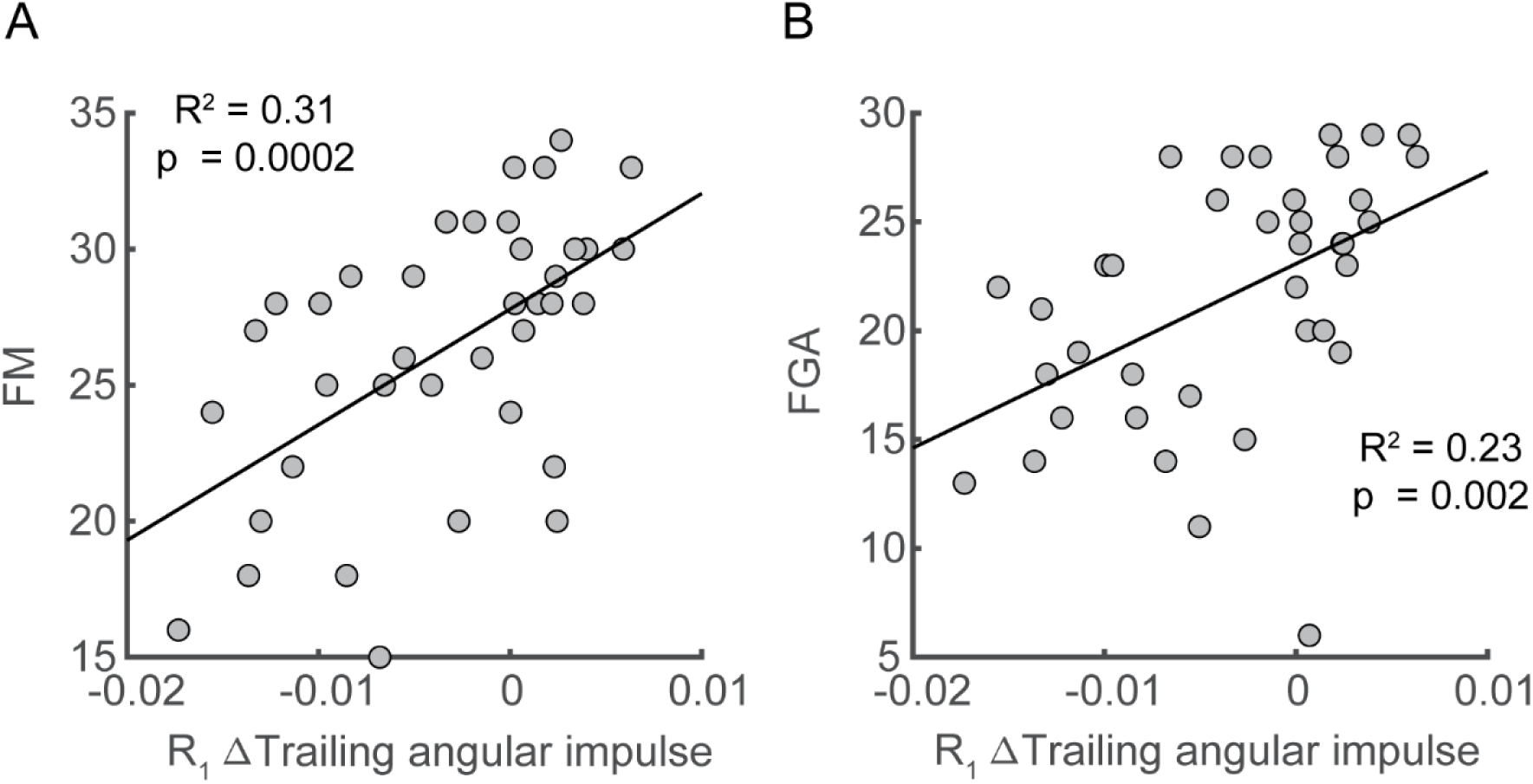
Associations between deviation in trailing angular impulse during the first recovery step from the pre-perturbation step in the sagittal plane and clinical assessments. Deviation of trailing angular impulse during the first recovery step from the pre-perturbation step was positively associated with (A) Fugl-Meyer score and (B) the Functional Gait Assessment only following paretic perturbations. FM: Fugl-Meyer; FGA: Functional Gait Assessment.

## 4 Discussion

The primary objective of this study was to determine how stroke affects the mechanical consequence of reactive control strategies in response to sudden treadmill accelerations. We found that perturbations to the paretic side led to more whole-body rotation during the perturbation step relative to non-paretic perturbations for people post-stroke and relative to neurotypical participants. To recover from these perturbations, neurotypical participants first used the perturbed stance limb to decrease the forward pitch impulse during the perturbed step. Then, during the double support phase of the first recovery step, they increased the leading limb angular impulse and decreased the forward trailing angular impulse by the perturbed limb relative to the pre-perturbation step. These reactive control strategies allowed neurotypical participants to restore the whole-body angular momentum to baseline levels within two steps.

In contrast to neurotypical participants, following paretic perturbations, people post-stroke did not decrease the forward angular impulse by the stance limb during the perturbed steps. People post-stroke also did not increase the leading limb angular impulse using the paretic leg or reduce the trailing limb angular impulse using their paretic leg during the double support phase of the first recovery step. However, when comparing responses to paretic versus non-paretic perturbations, we found that people post-stroke reduced their trailing limb angular impulse during the first recovery step more following non-paretic perturbations. Overall, people post-stroke primarily relied on their non-paretic limb to restore balance in contrast to neurotypical individuals who generated responses with substantial contributions from both limbs.

People post-stroke required more recovery steps to restore whole-body angular momentum than neurotypical individuals. Studies investigating postural control have used the number of recovery steps to quantify people’s ability to maintain balance in response to perturbation, and the use of multiple recovery steps is indicative of higher fall risk (Hilliard et al., 2008; Maki & McIlroy, 2006). For example, older adults, particularly those with a fall history, had a greater tendency to adopt multiple steps following a waist pull when standing compared to young adults (Mille et al., 2013). Additionally, people post-stroke needed more steps to restore balance following stance perturbations compared to age-matched controls (Martinez et al., 2019). Our results extended these observations to perturbations during walking by showing that people post-stroke needed one more recovery step following the treadmill-induced, slip-like perturbations to restore balance compared with age-matched neurotypical participants.

The increase in integrated whole-body angular momentum following paretic perturbations was higher than for non-paretic perturbations, indicating that people tended to fall forward more during paretic perturbations than non-paretic perturbations. The increase in the integrated whole-body angular momentum during the paretic perturbation step was about ~1.5 times higher than that on the non-paretic side. The increase in whole-body angular momentum from the pre-perturbation step following paretic perturbations was also higher than that for neurotypical participants, indicating that people post-stroke have impaired regulation of whole-body dynamics following paretic perturbations, which is in line with prior work indicating greater instability during backward losses of balance following paretic perturbations (Kajrolkar & Bhatt, 2016).

We observed marked differences in the mechanics of the most rapid balance correcting responses following paretic versus non-paretic perturbations. Neurotypical participants responded to the perturbations by modulating the stance limb ground reaction force toward the end of the perturbed steps to reduce forward angular impulse. However, at the group level, post-stroke participants did not reduce this impulse when the paretic limb was perturbed. This could be because the paretic perturbation steps were on average 145ms shorter than the non-paretic perturbation steps in our study. As a result, there might not be sufficient time for stroke participants to reduce the angular impulse during the stance phase during the paretic perturbation step. Additionally, people post-stroke may have delayed reactions to paretic perturbations due to sensory transmission or processing deficits, which would contribute to the increased forward loss of balance during paretic perturbations (Sharafi et al., 2016; C. Wutzke et al., 2013; C. J. Wutzke et al., 2015).

People post-stroke also showed impairments in the ability to increase leading limb angular impulse using the paretic limb relative to the non-paretic limb. Angular impulse is determined by the distance between the ground reaction force vectors the body’s center mass, reaction force magnitudes, and time duration of the forces applied. Neurotypical individuals regulate their ground reaction force vectors so that the vectors intersect slightly above the CoM throughout the gait cycle (Gruben & Boehm, 2012; Maus et al., 2010). This control strategy was also evident in neurotypical participants during perturbations that were generated by stepping down from a camouflaged curb (Vielemeyer et al., 2019). The ground reaction force vector from the leading limb continued to be directed above the CoM to generate a backward moment about the CoM to counteract the forward fall (Vielemeyer et al., 2019). However, this stabilization strategy of using ground reaction forces to control body dynamics may not be feasible in people post-stroke as the paretic limb may have limited ability to control force vector orientation relative to the center of mass compared to neurotypical participants (Boehm & Gruben, 2016; Rogers et al., 2004). Although people post-stroke can increase their step length to restore balance following a forward fall (Haarman et al., 2017), increasing step length may not be sufficient to change the leading limb angular impulse. We found no association between the increase in leading limb angular impulse during the first recovery step and the increase in step length or distance between foot placement and CoM (all p>0.05). Thus, generating sufficient leading limb angular impulse to arrest a forward loss of balance requires regulation of both ground reaction force and foot placement. Increasing paretic ground reaction force at the leading limb during the double support phase may generate high impact loading at the paretic limb and potentially cause knee collapse due to the weakness at the knee extensors. Thus, limiting the increase in leading limb impact angular impulse may be a protective mechanism for people post-stroke to avoid injury but additional study is needed to confirm this hypothesis.

Additionally, during the recovery step, people post-stroke did not reduce their paretic trailing limb angular impulse following the forward losses of balance to the same extent as they did with the non-paretic limb. At the beginning of the first recovery step, neurotypical participants reduced the forward angular impulse generated by the trailing limb, which likely limited the forward loss of balance caused by the sudden belt speed increase. One way to reduce the trailing limb forward angular impulse is by increasing the propulsive force in the anterior direction to generate a larger backward moment about the CoM if the moment arm is kept the same. The ability to increase the propulsive force requires the coordination of the hip flexor, knee extensor, and ankle plantarflexor moments of the trailing limb (Debelle et al., 2020; Pijnappels et al., 2005). Such a strategy may not be feasible for people post-stroke as they typically have abnormal coordination patterns which could prevent them from generating higher propulsive force at the trailing limb and redirect the ground reaction force vectors relative to the center of mass to reduce the overall angular impulse at the paretic limb (Allen et al., 2014; Finley et al., 2008; Hsiao et al., 2015; Sánchez et al., 2017).

Overall, our findings have important implications for interventions aimed at improving reactive balance control for people post-stroke. Specifically, our results may inform the design of perturbation-based interventions that seek to improve reactive stepping responses. The increased disturbance caused by paretic perturbations may reflect an inability to direct the ground reaction force vector of the paretic leg correctly relative to the body center of mass to reduce the forward loss of balance. Moreover, during the subsequent recovery steps following the perturbations, people post-stroke primarily relied on their non-paretic limb instead of coordinating both limbs to restore balance. Thus, future studies may investigate whether training could improve the ability to modulate paretic force vectors relative to the body center of mass so that momentum can be properly regulated throughout the gait cycle.

### 4.1 Limitations

In this current study protocol, we only elicited perturbations to induce forward loss of balance during walking with the same perturbation magnitudes for all participants. It remains to be determined if similar conclusions about the reactive stabilization strategies generated by people post-stroke extend to larger perturbations and perturbations in other directions. Moreover, although participants completed a familiarization trial to minimize the first trial effects, they may have adopted proactive active strategies that they would not typically employ due to heightened certainty about the likelihood of an upcoming perturbation.

## 5 Acknowledgments

We thank Cathy Broderick, Catherine Yunis, Ryan Novotny, and Sungwoo Park for their help with data collection. This work was supported by the Eunice Kennedy Shriver National Institute of Child Health & Human Development of the National Institutes of Health under Award Number R01HD091184 and SC CTSI (NIH/NCATS) through Grant UL1TR001855.

## 6 Author Contributions

C.L collected the data, analyzed data, and wrote the manuscript. J.L.M advised in data analysis and edited the manuscript. J.M.F designed the experiment, advised in data analysis, and edited the manuscript.

## 7 Conflict of Interest Statement

The authors declare that the research was conducted in the absence of any commercial or financial relationships that could be construed as a potential conflict of interest.

## Notes

### Competing Interest Statement

The authors have declared no competing interest.

## References

Adamczyk, P. G., & Kuo, A. D. (2009). Redirection of center-of-mass velocity during the step-to-step transition of human walking. Journal of Experimental Biology, 212(16), 2668–2678. https://doi.org/10.1242/jeb.027581

Allen, J. L., Kautz, S. A., & Neptune, R. R. (2014). Forward propulsion asymmetry is indicative of changes in plantarflexor coordination during walking in individuals with post-stroke hemiparesis. Clinical Biomechanics, 29(7), 780–786. https://doi.org/10.1016/j.clinbiomech.2014.06.001

Arene, N., & Hidler, J. (2009). Understanding motor impairment in the paretic lower limb after a stroke: A review of the literature. Topics in Stroke Rehabilitation, 16(5), 346–356. https://doi.org/10.1310/tsr1605-346

Berg, K. O., Maki, B. E., Williams, J. I., Holliday, P. J., & Wood-Dauphinee, S. L. (1992). Clinical and laboratory measures of postural balance in an elderly population. Archives of Physical Medicine and Rehabilitation, 73(11), 1073–1080. https://doi.org/0003-9993(92)90174-U [pii]

Boehm, W. L., & Gruben, K. G. (2016). Post-Stroke Walking Behaviors Consistent with Altered Ground Reaction Force Direction Control Advise New Approaches to Research and Therapy. Translational Stroke Research, 7(1), 3–11. https://doi.org/10.1007/s12975-015-0435-5

Buurke, T. J. W., Liu, C., Park, S., den Otter, R., & Finley, J. M. (2020). Maintaining sagittal plane balance compromises frontal plane balance during reactive stepping in people post-stroke. Clinical Biomechanics, 80, 105135. https://doi.org/10.1016/j.clinbiomech.2020.105135

Chen, G., Patten, C., Kothari, D. H., & Zajac, F. E. (2005). Gait differences between individuals with post-stroke hemiparesis and non-disabled controls at matched speeds. Gait & Posture, 22(1), 51–56. https://doi.org/10.1016/j.gaitpost.2004.06.009

Debelle, H., Harkness-Armstrong, C., Hadwin, K., Maganaris, C. N., & O’Brien, T. D. (2020). Recovery From a Forward Falling Slip: Measurement of Dynamic Stability and Strength Requirements Using a Split-Belt Instrumented Treadmill. Frontiers in Sports and Active Living, 2, 82. https://doi.org/10.3389/fspor.2020.00082

Dempster, W. T. (1955). Space requirements of the seated operator: Geometrical, kinematic, and mechanical aspects other body with special reference to the limbs. In WADC Technical Report (pp. 1–254). https://doi.org/AD087892

Dusane, S., Gangwani, R., Patel, P., & Bhatt, T. (2021). Does stroke-induced sensorimotor impairment and perturbation intensity affect gait-slip outcomes? Journal of Biomechanics, 118, 110255. https://doi.org/10.1016/j.jbiomech.2021.110255

Finley, J. M., Perreault, E. J., & Dhaher, Y. Y. (2008). Stretch reflex coupling between the hip and knee: Implications for impaired gait following stroke. Experimental Brain Research, 188(4), 529–540. https://doi.org/10.1007/s00221-008-1383-z

Fugl-Meyer, A. R., Jääskö, L., Leyman, I., Olsson, S., & Steglind, S. (1975). The post-stroke hemiplegic patient. 1. A method for evaluation of physical performance. Scandinavian Journal of Rehabilitation Medicine, 7(1), 13–31. https://doi.org/10.1038/35081184

Golyski, P. R., Vazquez, E., Leestma, J. K., & Sawicki, G. S. (2022). Onset timing of treadmill belt perturbations influences stability during walking. Journal of Biomechanics, 130, 110800. https://doi.org/10.1016/j.jbiomech.2021.110800

Gruben, K. G., & Boehm, W. L. (2012). Force direction pattern stabilizes sagittal plane mechanics of human walking. Human Movement Science, 31(3), 649–659. https://doi.org/10.1016/j.humov.2011.07.006

Haarman, J. A. M., Vlutters, M., Olde Keizer, R. A. C. M., Van Asseldonk, E. H. F., Buurke, J. H., Reenalda, J., Rietman, J. S., & Van Der Kooij, H. (2017). Paretic versus non-paretic stepping responses following pelvis perturbations in walking chronic-stage stroke survivors. Journal of NeuroEngineering and Rehabilitation, 14(1). https://doi.org/10.1186/s12984-017-0317-z

Hanavan, E. P. (1964). A Mathematical Model of the human body. Aerospace Medical Rsearch Laboratories, 1–149.

Havens, K. L., Mukherjee, T., & Finley, J. M. (2018). Analysis of biases in dynamic margins of stability introduced by the use of simplified center of mass estimates during walking and turning. Gait & Posture, 59(June 2017), 162–167. https://doi.org/10.1016/j.gaitpost.2017.10.002

Herr, H. M., & Popovic, M. (2008). Angular momentum in human walking. The Journal of Experimental Biology, 211(4), 467–481. https://doi.org/10.1242/jeb.008573

Higginson, J. S., Zajac, F. E., Neptune, R. R., Kautz, S. A., & Delp, S. L. (2006). Muscle contributions to support during gait in an individual with post-stroke hemiparesis. Journal of Biomechanics, 39, 1769–1777. https://doi.org/10.1016/j.jbiomech.2005.05.032

Hilliard, M. J., Martinez, K. M., Janssen, I., Edwards, B., Mille, M. L., Zhang, Y., & Rogers, M. W. (2008). Lateral Balance Factors Predict Future Falls in Community-Living Older Adults. Archives of Physical Medicine and Rehabilitation, 89(9), 1708–1713. https://doi.org/10.1016/j.apmr.2008.01.023

Honda, K., Sekiguchi, Y., Muraki, T., & Izumi, S. I. (2019). The differences in sagittal plane whole-body angular momentum during gait between patients with hemiparesis and healthy people. Journal of Biomechanics, 86, 204–209. https://doi.org/10.1016/j.jbiomech.2019.02.012

Hsiao, H., Knarr, B. A., Higginson, J. S., & Binder-Macleod, S. A. (2015). The relative contribution of ankle moment and trailing limb angle to propulsive force during gait. Human Movement Science, 39, 212–221. https://doi.org/10.1016/J.HUMOV.2014.11.008

Kajrolkar, T., & Bhatt, T. (2016). Falls-risk post-stroke: Examining contributions from paretic versus non paretic limbs to unexpected forward gait slips. Journal of Biomechanics, 49(13), 2702–2708. https://doi.org/10.1016/j.jbiomech.2016.06.005

Kajrolkar, T., Yang, F., Pai, Y.-C. Y.-C., & Bhatt, T. (2014). Dynamic stability and compensatory stepping responses during anterior gait-slip perturbations in people with chronic hemiparetic stroke. Journal of Biomechanics, 47(11), 2751–2758. https://doi.org/10.1016/j.jbiomech.2014.04.051

Kirker, S. G. B., Simpson, D. S., Jenner, J. R., & Wing, A. M. (2000). Stepping before standing: Hip muscle function in stepping and standing balance after stroke. J Neurol Neurosurg Psychiatry, 68, 458–464.

Kurz, M. J., Arpin, D. J., & Corr, B. (2012). Differences in the dynamic gait stability of children with cerebral palsy and typically developing children. Gait and Posture, 36(3), 600–604. https://doi.org/10.1016/j.gaitpost.2012.05.029

Lauzière, S., Miéville, C., Betschart, M., Aissaoui, R., & Nadeau, S. (2015). Plantarflexor weakness is a determinant of kinetic asymmetry during gait in post-stroke individuals walking with high levels of effort. Clinical Biomechanics, 30(9), 946–952. https://doi.org/10.1016/j.clinbiomech.2015.07.004

Leddy, A. L., Crowner, B. E., & Earhart, G. M. (2011). Functional gait assessment and balance evaluation system test: Reliability, validity, sensitivity, and specificity for identifying individuals with Parkinson disease who fall. Physical Therapy, 91(1), 102–113. https://doi.org/10.2522/ptj.20100113

Liu, C., De Macedo, L., & Finley, M. J. (2018). Conservation of Reactive Stabilization Strategies in the Presence of Step Length Asymmetries during Walking. Frontiers in Human Neuroscience, 12, 251. https://doi.org/doi:10.3389/fnhum.2018.00251

Maki, B. E., & McIlroy, W. E. (2006). Control of rapid limb movements for balance recovery: Age-related changes and implications for fall prevention. Age and Ageing, 35(SUPPL.2). https://doi.org/10.1093/ageing/afl078

Marigold, D. S., Eng, J. J., & Timothy Inglis, J. (2004). Modulation of ankle muscle postural reflexes in stroke: Influence of weight-bearing load. Clinical Neurophysiology, 115(12), 2789–2797. https://doi.org/10.1016/j.clinph.2004.07.002

Martelli, D., Monaco, V., Bassi Luciani, L., Micera, S., Luciani, L. B., & Micera, S. (2013). Angular Momentum During Unexpected Multidirectional Perturbations Delivered While Walking. IEEE Transactions on Biomedical Engineering, 60(7), 1785–1795. https://doi.org/10.1109/TBME.2013.2241434

Martinez, K. M., Rogers, M. W., Blackinton, M. T., Samuel Cheng, M., & Mille, M. L. (2019). Perturbation-Induced Stepping Post-stroke: A pilot study demonstrating altered strategies of both legs. Frontiers in Neurology, 10(JUL). https://doi.org/10.3389/fneur.2019.00711

Mathiyakom, W., & McNitt-Gray, J. L. (2008). Regulation of angular impulse during fall recovery. In Journal of Rehabilitation Research and Development. https://doi.org/10.1682/JRRD.2008.02.0033

Maus, H. M., Lipfert, S. W., Gross, M., Rummel, J., & Seyfarth, A. (2010). Upright human gait did not provide a major mechanical challenge for our ancestors. Nature Communications, 1(6). https://doi.org/10.1038/ncomms1073

Mille, M. L., Johnson-Hilliard, M., Martinez, K. M., Zhang, Y., Edwards, B. J., & Rogers, M. W. (2013). One step, two steps, three steps more… Directional vulnerability to falls in community-dwelling older people. Journals of Gerontology - Series A Biological Sciences and Medical Sciences, 68(12 A), 1540–1548. https://doi.org/10.1093/gerona/glt062

Nott, C. R., Neptune, R. R., & Kautz, S. A. (2014). Relationships between frontal-plane angular momentum and clinical balance measures during post-stroke hemiparetic walking. Gait & Posture, 39(1), 129–134. https://doi.org/doi:10.1016/j.gaitpost.2013.06.008.

Olney, S. J., & Richards, C. (1996). Hemiparetic gait following stroke. Part 1: Characteristics. Gait & Posture, 4, 136–148. https://doi.org/10.1016/0966-6362(96)01063-6

Park, S., Liu, C., Sánchez, N., Tilson, J. K., Mulroy, S. J., & Finley, J. M. (2021). Using Biofeedback to Reduce Step Length Asymmetry Impairs Dynamic Balance in People Poststroke. Neurorehabilitation and Neural Repair, 35(8), 738–749. https://doi.org/10.1177/15459683211019346

Patel, P. J., & Bhatt, T. (2017). Fall risk during opposing stance perturbations among healthy adults and chronic stroke survivors. Experimental Brain Research, 1(236), 619–628. https://doi.org/10.1007/s00221-017-5138-6

Pijnappels, M., Bobbert, M. F., & Dieën, J. H. van. (2005). Push-off reactions in recovery after tripping discriminate young subjects, older non-fallers and older fallers. Gait & Posture, 21(4), 388–394. https://doi.org/10.1016/j.gaitpost.2004.04.009

Potocanac, Z., de Bruin, J., van der Veen, S., Verschueren, S., van Dieën, J., Duysens, J., & Pijnappels, M. (2014). Fast online corrections of tripping responses. Experimental Brain Research, 232(11), 3579–3590. https://doi.org/10.1007/s00221-014-4038-2

Powell, L. E., & Myers, A. M. (1995). The Activities-specific Balance Confidence (ABC) Scale. The Journals of Gerontology: Series A: Biological Sciences and Medical Sciences, 50(1), M28–M34. https://doi.org/10.1093/gerona/50A.1.M28

Reisman, D. S., Wityk, R., Silver, K., Bastian, A. J., & Manuscript, A. (2009). Split-Belt Treadmill Adaptation Transfers to Overground Walking in Persons Poststroke. Physical Therapy, 23(7), 735–744. https://doi.org/10.1177/1545968309332880.Split-Belt

Roeles, S., Rowe, P. J., Bruijn, S. M., Childs, C. R., Tarfali, G. D., Steenbrink, F., & Pijnappels, M. (2018). Gait stability in response to platform, belt, and sensory perturbations in young and older adults. Medical & Biological Engineering & Computing, 56(12), 2325–2335. https://doi.org/10.1007/s11517-018-1855-7

Roerdink, M., Geurts, A. C. H., de Haart, M., & Beek, P. J. (2009). On the Relative Contribution of the Paretic Leg to the Control of Posture After Stroke. Neurorehabilitation and Neural Repair, 23(3), 267–274. https://doi.org/10.1177/1545968308323928

Rogers, L. M., Brown, D. A., & Gruben, K. G. (2004). Foot force direction control during leg pushes against fixed and moving pedals in persons post-stroke. Gait and Posture, 19(1), 58–68. https://doi.org/10.1016/S0966-6362(03)00009-2

Rybar, M. M., Walker, E. R., Kuhnen, H. R., Ouellette, D. R., Berrios, R., Hunter, S. K., & Hyngstrom, A. S. (2014). The stroke-related effects of hip flexion fatigue on over ground walking. Gait and Posture, 39(4), 1103–1108. https://doi.org/10.1016/j.gaitpost.2014.01.012

Salot, P., Patel, P., & Bhatt, T. (2016). Reactive Balance in Individuals With Chronic Stroke: Biomechanical Factors Related to Perturbation-Induced Backward Falling. Physical Therapy, 96(3), 338–347. https://doi.org/10.2522/ptj.20150197

Sánchez, N., Acosta, A. M., Lopez-Rosado, R., Stienen, A. H. A.;, & Dewald, J. P. A.; (2017). Lower Extremity Motor Impairments in Ambulatory Chronic Hemiparetic Stroke: Evidence for Lower Extremity Weakness and Abnormal Muscle and Joint Torque Coupling Patterns. Neurorehabilitation and Neural Repair, 31(9), 814–826. https://doi.org/10.1177/1545968317721974

Schmid, A. A., Yaggi, H. K., Burrus, N., McClain, V., Austin, C., Ferguson, J., Fragoso, C., Sico, J. J., Miech, E. J., Matthias, M. S., Williams, L. S., & Bravata, D. M. (2013). Circumstances and consequences of falls among people with chronic stroke. Journal of Rehabilitation Research and Development, 50(9), 1277–1286. https://doi.org/10.1682/JRRD.2012.11.0215

Sharafi, B., Hoffmann, G., Tan, A. Q., & Y. Dhaher, Y. (2016). Evidence of impaired neuromuscular responses in the support leg to a destabilizing swing phase perturbation in hemiparetic gait. Experimental Brain Research, 234(12), 3497–3508. https://doi.org/10.1007/s00221-016-4743-0

Silverman, A. K., & Neptune, R. R. (2011). Differences in whole-body angular momentum between below-knee amputees and non-amputees across walking speeds. Journal of Biomechanics, 44(3), 379–385. https://doi.org/10.1016/j.jbiomech.2010.10.027

Sloot, L. H., Van Den Noort, J. C., Van Der Krogt, M. M., Bruijn, S. M., & Harlaar, J. (2015). Can treadmill perturbations evoke stretch reflexes in the calf muscles? PLoS ONE, 10(12). https://doi.org/10.1371/journal.pone.0144815

Song, J., Sigward, S., Fisher, B., & Salem, G. J. (2012). Altered dynamic postural control during step turning in persons with early-stage Parkinson’s disease. Parkinson’s Disease, 2012, 386962. https://doi.org/10.1155/2012/386962

Vielemeyer, J., Grießbach, E., & Müller, R. (2019). Ground reaction forces intersect above the center of mass even when walking down visible and camouflaged curbs. Journal of Experimental Biology, 222(14). https://doi.org/10.1242/jeb.204305

Vlutters, M., Van Asseldonk, E. H. F., & Van Der Kooij, H. (2016). Center of mass velocity-based predictions in balance recovery following pelvis perturbations during human walking. Journal of Experimental Biology, 219(10), 1514–1523. https://doi.org/10.1242/jeb.129338

Weerdesteyn, V., De Niet, M., van Duijnhoven, H. J. R. R., Geurts, A. C. H. H., Niet, M. de, van Duijnhoven, H. J. R. R., & Geurts, A. C. H. H. (2008). Falls in individuals with stroke. The Journal of Rehabilitation Research and Development, 45(8), 1195. https://doi.org/10.1682/JRRD.2007.09.0145

Winter, D. A. (2009). Biomechanics and Motor Control of Human Movement. In Motor Control (Vol. 2nd). https://doi.org/10.1002/9780470549148

Wutzke, C. J., Faldowski, R. A., & Lewek, M. D. (2015). Individuals Poststroke Do Not Perceive Their Spatiotemporal Gait Asymmetries as Abnormal. Physical Therapy, 95(9), 1244–1253. https://doi.org/10.2522/ptj.20140482

Wutzke, C., Mercer, V., & Lewek, M. (2013). Influence of lower extremity sensory function on locomotor adaptation following stroke: A review. Topics in Stroke Rehabilitation, 20(3), 233–240. https://doi.org/10.1310/tsr2003-233

