## Supplementary material for "Impairments in the mechanical effectiveness of reactive balance control strategies during walking in people post-stroke"

### 1 Supplementary Results

#### 1.1 Whole-body angular momentum

We examined the effects of perturbations on the magnitude of integrated whole-body angular momentum for control participants and found a significant main effect of Step Type on angular momentum (F(1,144) = 34.9, p < 0.0001). Participants increased their forward rotation, indicated by a more negative *L_int_,* during the perturbation step relative to the pre-perturbation step (t(144) = -4.82, p < 0.0001). They then countered the effects of the perturbation during the first recovery step (R_1_) as indicated by a more positive *L_int_*  (t(144) = 4.7, p < 0.0001). Neurotypical participants restored whole-body angular momentum to levels comparable to those observed during the pre-perturbation step by the second recovery step (R_2_) (p = 0.86).

Similarly, we quantified the effects of perturbations on integrated angular momentum across the step cycle to for stroke participants. We found a main effect of Step Type (F(5,444) = 64.7, p <0.001) and an interaction between Leg and Step Type (F(5, 444) = 3.0, p = 0.011), but there was no main effect of Leg on the integrated angular momentum (F(1, 444) = 2.12, p = 0.14) . *L_int_* differed from the pre-perturbation step during the perturbation (PTB) step (t(444) = -6.96, p < 0.001), first recovery step (R_1_) (t(444) = 9.4, p < 0.001), and second recovery step (R_2_) (t(444) = -2.02, p = 0.04). There was no significant difference between $L_{int}$during the third recovery step and the pre-perturbation step (t(444) = -1.05, p = 0.29). Thus, people post-stroke generally restored their angular momentum to pre-perturbation values by the third recovery step. Additionally, 36 out of 38 stroke participants had larger increases in *L_int_* during the paretic perturbation step than the non-paretic perturbation step (t(444) = -2.3, p = 0.021), indicating that people post-stroke fell forward more when the perturbation occurred during paretic stance.

#### 1.2 Pitch impulse (ΔL_Stance_)

During the pre-perturbation step, the ground reaction force produced by the stance limb from midstance to the subsequent foot strike typically produced a forward pitch impulse for both neurotypical individuals and people post-stroke (Figure S1). There was no difference in the magnitude of the pitch impulse during the pre-perturbation step between stroke and neurotypical participants.

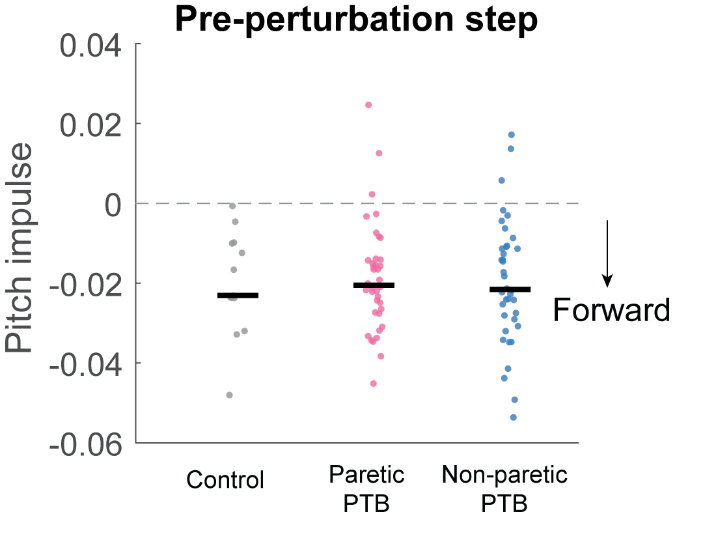

Figure S1. Median pitch impulse during the pre-perturbation step for control participants (Gray, N = 13) and during paretic (Pink) and non-paretic (Blue) steps for stroke participants (N=38). Each dot represents one participant. Black horizontal lines indicate the median across participants. Negative values indicate forward pitch impulse.

#### 1.3 Net angular impulse (ΔL_Net_)

We computed net angular impulse as the sum of leading limb angular impulse and trailing limb angular impulse. During the double support phase of the pre-perturbation step, net angular impulse was typically positive for neurotypical individuals (Figure S2A). During the perturbation step, neurotypical participants decreased the net angular impulse relative to the pre-perturbation step (t(48) = -4.7, p < 0.0001), which was consistent with the increase in forward momentum caused by the perturbation (Figure S2B).

For stroke participants, the net angular impulse during pre-perturbation steps was more negative during non-paretic steps than during paretic steps (t(444) = -4.4, p < 0.0001, Figure S2A). This indicates that people post-stroke arrested more of the forward momentum of the body during paretic versus non-paretic steps. During the perturbation step, stroke participants decreased the net angular impulse following both paretic (t(444) = -2.3, p = 0.018) and non-paretic perturbations (t(444) = -2.1, p = 0.034), which is consistent with a forward loss of balance (Figure S2B).

#### 1.4 Leading limb angular impulse (ΔL_Leading_)

During the pre-perturbation step, the ground reaction force generated by the leading leg of neurotypical control participants produced a backward angular impulse about the CoM during the double support phase (Figure S2C). The perturbations led to a reduction in backward impulse during the perturbation step compared to the pre-perturbation step (t(48) = -4.6, p < 0.0001, Figure S2D) and thus participants tended to fall forward during the perturbations.

For stroke participants, the leading limb angular impulse generated by the non-paretic leg was larger than that generated by the paretic leg (p < 0.0001) during the pre-perturbation step (Figure S2C). Similar to neurotypical participants, the leading limb angular impulse for stroke participants decreased from the pre-perturbation step during paretic (t(444) = -2.2, p = 0.027) and non-paretic perturbations (t(444) = -2.3, p = 0.02, Figure S2D).

#### 1.5 Trailing limb angular impulse (ΔL_Trailing_ )

During the pre-perturbation step, the ground reaction force by the trailing limb typically generated a forward moment about the body’s CoM, and thus the trailing limb angular impulse was negative for neurotypical participants (Figure S2E). There was no change in trailing limb angular impulse during the perturbation step from that measured during the pre-perturbation step (t(48) = 0.17, p = 0.86, Figure S2F).

For stroke participants during the pre-perturbation step, the magnitude of trailing limb impulse generated by the non-paretic trailing leg was higher than that generated by the paretic leg (t(444) = 7.5, p < 0.0001) and higher than that generated by the neurotypical participants (U = 472, Bonferroni corrected p = 0.012; Figure S2E). Similar to neurotypical participants, there were no changes in trailing limb angular impulse from the pre-perturbation step for either paretic (t(444) = 0.50, p = 0.62) or non-paretic perturbations (t(444) = 0.41, p = 0.68, Figure S2F).

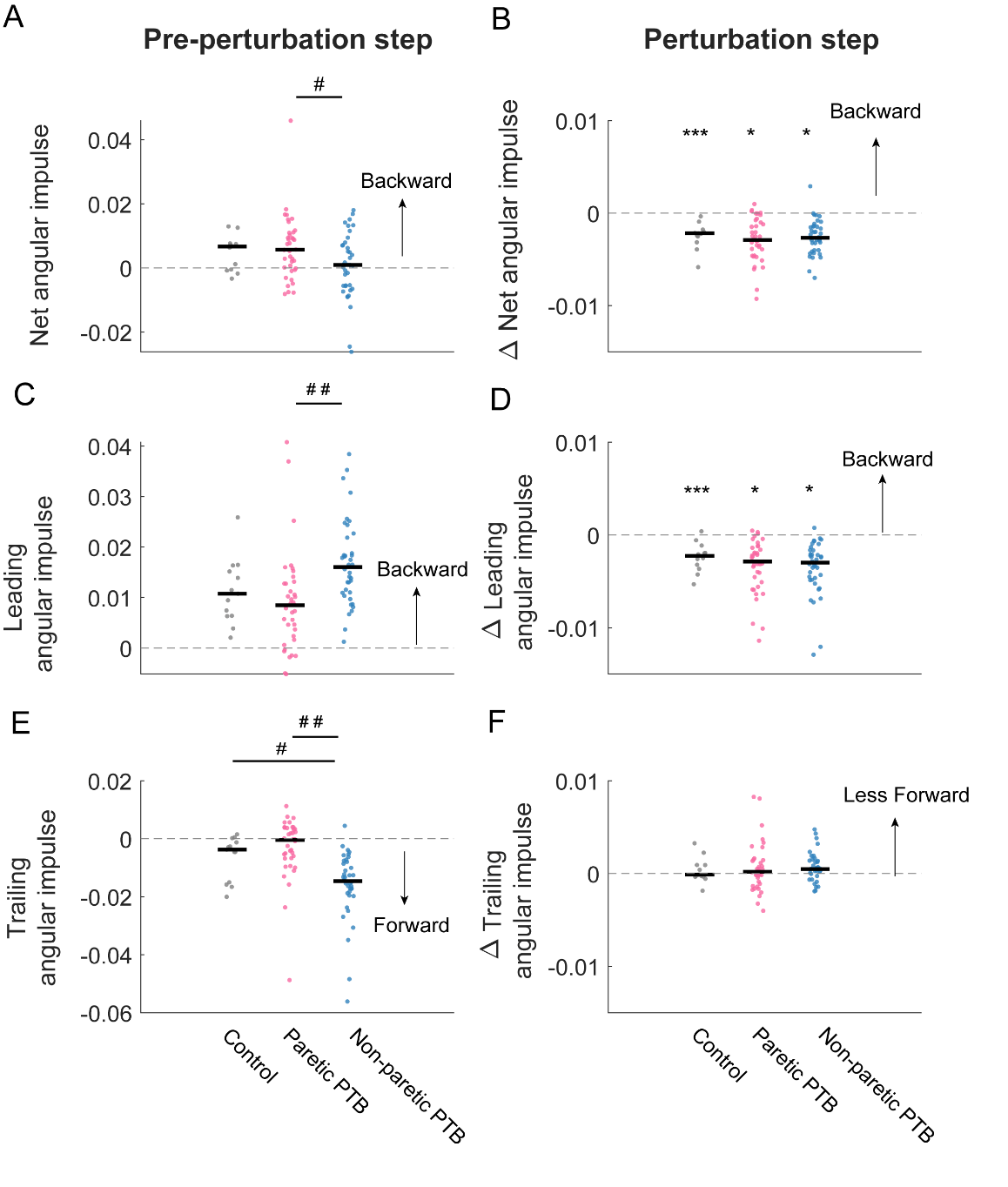

Figure S2. Median net, leading limb, and trailing limb angular impulse_,_ during the pre-perturbation and perturbation step. The left column shows median net (A), leading limb (C), and trailing limb (E)angular impulse during the pre-perturbation step across control participants (Gray, N = 13) and stroke participants (N=38) during paretic (Pink) and non-paretic (Blue) steps. The right column shows median changes in net (B),leading limb (D), trailing limb angular impulse,(E) during the perturbation stes compared to those measured during the pre-perturbation step. Each dot represents one participant. Black horizontal lines indicate the median across participants. The asterisks (*)indicate whether values were statistically different from zero (*p<0.05, **p<0.001, ***p<0.0001). The hashes (#) indicate when comparisons between groups are significantly different (#p<0.05, ##p<0.001).

**Table S1:** Statistical results from the linear mixed effect models examining changes in each outcome during the perturbation step (PTB) and the first recovery step (R_1_) relative to the pre-perturbation step (Pre-PTB) for perturbations to neurotypical participants and perturbations to non-paretic and paretic side for stroke participants.

|  | **Outcome Measures** | **Step versus Pre-PTB** | **Perturbation** | **DF** | **t value** | **P value** |
| --- | --- | --- | --- | --- | --- | --- |
| 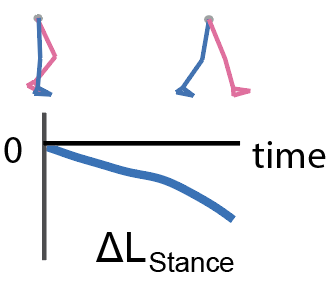 | **Pitch impulse** | **PTB** | Control | 48 | 3.7 | **0.0005** |
|  | ΔL_stance_ |  | Non-paretic | 444 | 2.8 | **0.005** |
|  |  |  | Paretic | 444 | 0.4 | 0.67 |
| 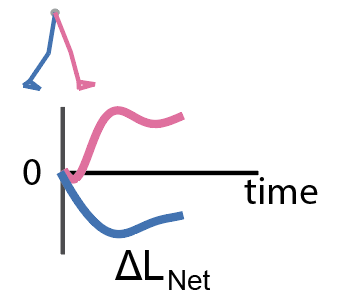 |  | **PTB** | Control | 48 | -4.7 | **<0.0001** |
|  |  |  | Non-paretic | 444 | -2.1 | **0.034** |
|  | **Net angular impulse**  ΔL_Net_  = ΔL_Leading_ + ΔL_Trailing_ |  | Paretic | 444 | -2.3 | **0.018** |
|  |  | **R_1_** | Control | 48 | 12.1 | **<0.0001** |
|  |  |  | Non-paretic | 444 | 3.6 | **0.0003** |
|  |  |  | Paretic | 444 | 1.64 | 0.1 |
| 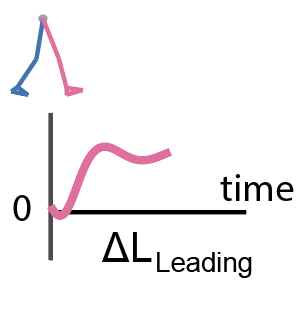 |  | **PTB** | Control | 48 | -4.6 | **<0.0001** |
|  |  |  | Non-paretic | 444 | -2.2 | **0.028** |
|  | **Leading angular impulse**  ΔL_Leading_ |  | Paretic | 444 | -2.3 | **0.02** |
|  |  | **R_1_** | Control | 48 | 5.7 | **<0.0001** |
|  |  |  | Non-paretic | 444 | 1.8 | 0.075 |
|  |  |  | Paretic | 444 | 3.5 | **0.0006** |
| 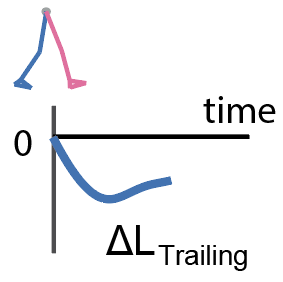 |  | **PTB** | Control | 48 | 0.17 | 0.86 |
|  |  |  | Non-paretic | 444 | 0.41 | 0.68 |
|  | **Trailing angular impulse**  ΔL_Trailing_ |  | Paretic | 444 | 0.5 | 0.62 |
|  |  | **R_1_** | Control | 48 | 5.1 | **<0.0001** |
|  |  |  | Non-paretic | 444 | 1.1 | 0.27 |
|  |  |  | Paretic | 444 | -2.0 | **0.047** |

**Table S2**: Statistical analysis for comparing changes in each outcome measures during the perturbation step (PTB) and the first recovery step (R_1_) relative to the pre-perturbation step (Pre-PTB) between control participants and stroke participants and between non-paretic and paretic perturbations.

|  | **Step versus  Pre-PTB** | **Control vs. Paretic** | **Control vs. Non-paretic** | **Non-paretic vs. Paretic** |
| --- | --- | --- | --- | --- |
| **Pitch impulse** |  |  |  |  |
| ΔL_stance_ | **PTB** | P = 0.78 | P = 0.1 | P = 0.088 |
| **Net angular impulse** | **PTB** | P = 1 | P = 1 | P = 0.87 |
| ΔL_Net_ | **R_1_** | P = 0.051 | **P = 0.0072** | P = 0.17 |
| **Leading angular impulse** | **PTB** | P = 1 | P = 0.72 | P = 0.93 |
| ΔL_Leading_ | **R_1_** | P = 1 | P = 0.42 | P = 0.24 |
| **Trailing angular impulse** | **PTB** | P = 1 | P = 0.78 | P = 0.95 |
| ΔL_Trailing_ | **R_1_** | **P = 0.0033** | P = 0.69 | **P = 0.029** |
